# The NALCN regulator UNC-80 functions in a subset of interneurons to affect *C. elegans* avoidance response to the plant hormone methyl salicylate

**DOI:** 10.1101/677070

**Authors:** Chuanman Zhou, Jintao Luo, Xiaohui He, Qian Zhou, Yunxia He, Xiaoqin Wang, Long Ma

## Abstract

NALCN (<u>Na</u>^+^ leak <u>c</u>hannel, <u>n</u>on-selective), UNC80 and UNC79 form a non-selective, voltage-independent cation channel complex that affects a broad array of neuronal activities. The molecular and neuronal mechanisms underlying the functions of the NALCN complex remain unclear. In a screen for *Caenorhabditis elegans* mutants with defective avoidance response to the plant hormone methyl salicylate (MeSa), we isolated novel loss-of-function (lf) mutations in *unc-80* and *unc-79*. *unc-80* and *unc-79* lf mutants exhibited defective MeSa avoidance but wild type-like responses to other odorants. Lf mutants of *C. elegans nca/NALCN* exhibited similar MeSa-specific avoidance defect, while lf mutants of the NALCN regulatory gene *nlf-1* avoided MeSa like wild type. Using fluorescent transgenic animals, we identified a subset of *unc-80*-expressing neurons. Neuron-specific transgene rescue and knockdown experiments suggest that a subset of interneurons, primarily including AVA, AVE and AVG, might play a necessary and sufficient role in mediating *unc-80* regulation of the MeSa avoidance. We found that *unc-79* was expressed in neurons largely overlapping those expressing *unc-80*, which is supported by the rescue of *unc-80(lf)* defects using an *unc-80* transgene driven by an *unc-79* promoter. We also suggest that *C. elegans* locomotion responds more sensitively to the changes of expression levels of *NALCN*-related genes than the MeSa avoidance does. Together, our results identified NALCN-related genes as key regulators of the MeSa avoidance behavior and provided novel genetic and neuronal insights into the function of the NALCN channel complex.

**Author summary:** NALCN (<u>Na</u>^+^ leak <u>c</u>hannel, <u>n</u>on-selective) is a non-selective, voltage-independent cation channel that affects multiple neuronal activities and behaviors. Mutations in NALCN and its regulator UNC80 can cause serious neurological diseases. The regulation and function of the NALCN channel complex remain to be understood. From a genetic screen, we surprisingly found that the nematode *Caenorhabditis elegans* requires NALCN and its two regulators UNC-80 and UNC-79 to escape from the plant stress hormone methyl salicylate (MeSa). Using methods including transgenic neuronal labeling, rescues and knockdowns, we found that *unc-80*-expression in a subset of head interneurons, including AVA, AVE and AVG, might be necessary and sufficient to elicit the MeSa avoidance response. We also found that *unc-79* functions in overlapping neurons as *unc-80* to regulate *C. elegans* behaviors. Our findings provide novel molecular and neuronal mechanisms for understanding the regulation and function of the NALCN channel complex.

## Introduction

The NALCN (<u>Na</u>^+^ leak <u>c</u>hannel, <u>n</u>on-selective) channel is a non-selective, voltage-independent cation channel broadly expressed in the animal kingdom [1, 2]. NALCN functions in neurons to balance the K^+^ leak, set the resting membrane potential, regulate spontaneous firing of neurons and modulate membrane potential in response to environmental stimuli [2].

NALCN affects a variety of biological processes in mammals. NALCN mutant mice die within 24 hrs after birth due to disrupted respiratory rhythm [3]. The channel has been implicated in the regulation of pacemaking activity in mouse gastrointestinal tract [4], clock neuron rhythms [5], firing and glycolytic sensitivity of substantia nigra pars reticulata neurons [6], excitability of the retrotrapezoid nucleus neurons [7], rapid eye movement sleep [8], and rhythmic stability within the respiratory network [9]. Mutations in NALCN and its regulatory protein UNC80 cause human diseases [10–15] that are collectively called NALCN channelopathies [11].

NALCN also regulates neuronal activities and behaviors in invertebrates. *Drosophila NALCN* mutants exhibit the narrow abdomen *(na)* phenotype, disrupted circadian rhythm and resistance to the volatile anesthetics halothane [16–18]. *C. elegans* has two *NALCN* homologs, *nca-1* and *nca-2*, that function redundantly to regulate the response to volatile anesthetics [19], recycling of synaptic vesicles [20], neural circuit activity [21], motor behavior [22], the propagation of neuronal activity from cell bodies to synapses [23], ethanol responses [24] and developmentally timed sleep [25].

In mammalian cells, NALCN channel can be regulated by G protein-coupled receptor TACR1, Src kinases [26], a [Ca^2+^]-sensing G protein-coupled receptor [27] and the M3 muscarinic receptors (M3R) [28]. In *C. elegans*, NCA channels function downstream of the Gq-Rho pathway [29], can be negatively regulated by dopamine through the D2-like dopamine receptor DOP-3 [30] and may interact with the SEK-1 p38 MAPK pathway [31].

Studies in *C. elegans*, *Drosophila* and mice identified the conserved proteins UNC79 and UNC80 (orthologs of *C. elegans* UNC-79 and UNC-80, respectively) as key regulators the NALCN channel [19,20,22–24,26,27,32–34]. In *C. elegans*, loss-of-function (lf) mutants of *unc-79* or *unc-80* phenocopy *nca-2(lf); nca-1(lf)* double mutants [19,20,22,23,25]. At the molecular level, *unc-79* and *unc-80* are required for the proper expression and axonal localization of NCA channels [19,20,23]. In *Drosophila*, lf mutations in *unc79*, *unc80* or *NALCN* cause indistinguishable defects in circadian locomotion rhythmicity [32] and similarly abnormal responses to halothane [19]. In mice, UNC79 and UNC80 are required for NALCN channel activity [26, 27]. Recently, an ER-associated protein, NLF-1, was found to promote axonal localization of NCA channels in *C. elegans* [35]. The *Drosophila* ortholog of NLF-1 is required for the regulation of circadian neuron excitability [5].

We previously found that *C. elegans* exhibits a strong avoidance response to the plant hormone methyl salicylate (MeSa) and multiple neuronal genes can affect this behavior [36]. To identify new genes involved, we screened for mutants with defective MeSa avoidance. Surprisingly, the screen isolated novel lf mutations in *unc-80* and *unc-79*. In this study, we examined how *nca, unc-79*, *unc-80* and *nlf-1* affect *C. elegans* avoidance to MeSa and locomotion. We also identified a subset of neurons that expresses *unc-80* and analyzed the roles of these neurons in mediating *unc-80* functions. Our findings provide novel genetic and neuronal mechanisms for understanding the regulation and function of the NALCN channel complex.

## Results

### A screen identified mutants with defective MeSa avoidance response

To identify novel genes required for the MeSa avoidance behavior, we performed a genetic screen for mutants that failed to avoid MeSa. The screen isolated 15 lines (Fig. S1A). Genetic complementation tests divided the mutations to three groups (Table S1).

To investigate whether the mutants had defects in avoiding MeSa specifically or sensing odorants in general, we examined a representative in each complementation group for chemotaxis (Table S1, *mac379*, *mac383* and *mac387*, respectively). *mac379* and *mac383* mutants exhibited wild type-like responses (Fig. S1B) to repelling odorants 1-octanol and 2-nonanone, and attractive odorants diacetyl and benzaldehyde [37], suggesting that these mutants might carry MeSa-specific defects. The third mutant, *mac387*, was defective in detecting each odorant (Fig. S1B). We postulate that *mac387* might cause broad defects in chemotaxis and did not further analyze it.

### Loss-of-function mutations in *unc-80* and *unc-79* caused MeSa avoidance defect

To identify the causal mutations leading to the MeSa avoidance defect, we determined the genomic sequences of *mac379*, *mac382* and *mac388* mutants from group 1 and *mac383* from group 2 (Table S1). A comparison of candidate genes found that *unc-80* was the only affected gene shared by *mac379*, *mac382* and *mac388*, with W1524stop, W220stop and W1967stop as the respective nonsense mutations (Table S1). Besides W1524stop, *mac379* also carried a missense mutation (G927R) in *unc-80* (Fig. S2A and Table S1). Sanger sequencing identified distinct nonsense or splice site mutations in *unc-80* from other mutants of group 1 (Table S1).

Meanwhile, we identified an R885stop mutation in *unc-79* among the candidate genes for *mac383* (group 2) (Table S1 and Fig. S2B). Considering that *unc-80* and *unc-79* interact with *nca* to affect *C. elegans* behavior [19,20,23], we speculated that *unc-79* might be the causal gene of *mac383*. Indeed, Sanger sequencing identified distinct deletion/frameshift mutations in *unc-79* from *mac389* and *mac393* mutants of group 2 (Table S1).

To verify that *unc-79* is required for the MeSa avoidance behavior, we performed transgene rescue experiments. Driven by a long endogenous promoter (Fig. 1A, *P_L_unc-79a*, 5.0 kb), an *unc-79a* gDNA transgene (Fig. 1A) strongly rescued the defective locomotion and MeSa avoidance of *unc-79(mac383)* mutants (Fig. 1B). Driven by a short promoter (Fig. 1A, *P_S_unc-79a*, 2.0 kb), the *unc-79a* gDNA transgene only recued the defective MeSa avoidance but not the defective locomotion (Fig. 1B). We also examined a shorter *unc-79* isoform, *unc-79b* (Fig. S2B and 1A, *Punc-79b*), and found that it failed to rescue either defective behavior (Fig. 1B).

**Figure 1.**
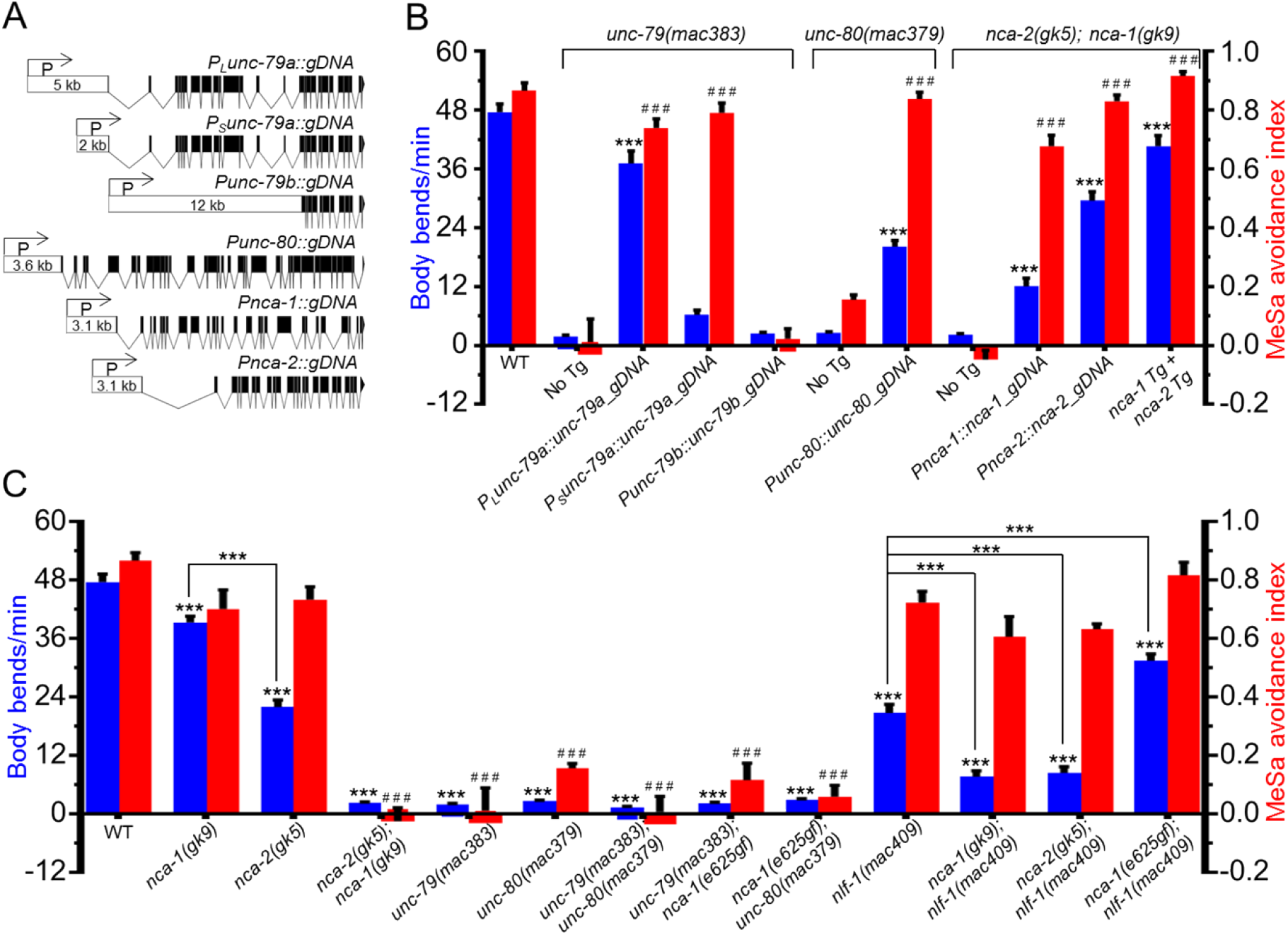
The effects of *unc-79*, *unc-80* and *nca* mutations on MeSa avoidance and locomotion. (A) Transgene structures. The length of each promoter is indicated. (B) Transgene rescue of the defective locomotion and the defective MeSa avoidance of *unc-79*, *unc-80* and *nca* lf mutants. The locomotion (body bends/min, blue columns) and MeSa avoidance indexes (red columns) were based on two independent transgenic lines. For locomotion, 20 animals were assayed for each line with 40 animals assayed in total. For MeSa avoidance indexes, each dataset was based on three biological replicates for each line with six in total. Statistics: Student’s *t*-test. ***, p<0.001. Error bars: standard error of mean. Comparisons were made between transgenic lines and their respective lf mutants. (C) The locomotion (body bends/min, blue columns, 40 animals per strain) and MeSa avoidance indexes (red columns, three to six biological replicates) of single and double mutants. Genotypes are labeled at bottom. Statistics: Bonferroni multiple comparison with one-way ANOVA. *** or ^###^, p<0.001. Error bars: standard error of mean. Note: data for wildtype, *unc-79(lf)*, *unc-80(lf)* and *nca(lf)* animals were repeatedly used for comparison purpose.

Similarly, an *unc-80* gDNA transgene that covers all annotated *unc-80* isoforms (Fig. S2A) driven by an *unc-80* endogenous promoter (Fig. 1A, *Punc-80*, 3.6 kb) can significantly rescue the defective locomotion and MeSa avoidance of *unc-80(mac379)* mutants (Fig. 1B). Compared to the completely rescued MeSa avoidance, the defective locomotion was only partially rescued (Fig. 1B).

*unc-79* and *unc-80* lf mutants are hypersensitive to the anesthetic halothane and exhibit the “fainter” phenotype [19,20,23,38–41]. Similar “fainter” phenotype was observed in all *unc-80* and *unc-79* mutants isolated in this study (unpublished observation). In addition, *unc-79(e1291)* and *unc-80(e1272)* animals, two previously described lf mutants [20], exhibited defective MeSa avoidance resembling that of *unc-79(mac383)* or *unc-83(mac379)* mutants (unpublished observation). Together, these results suggest that *unc-79* and *unc-80* are required for *C. elegans* MeSa avoidance behavior and we isolated lf mutations in *unc-79* and *unc-80*.

### *nca-1* and *nca-2* were redundantly required for the MeSa avoidance response

*C. elegans nca-1* and *nca-2* encode functionally redundant NALCN channels [19,20,23,40]. To investigate whether *nca* is required for the MeSa avoidance behavior, we examined *nca* single mutants and double mutants. We found that either *nca-1(gk9lf)* or *nca-2(gk5lf)* single mutants exhibited wild type-like MeSa avoidance (Fig. 1C), while *nca-2(lf); nca-1(lf)* double mutants were strongly defective in avoiding MeSa (Fig. 1B and 1C). An *nca-1* gDNA transgene (Fig. 1A, *Pnca-1*), an *nca-2* gDNA transgene (Fig. 1A, *Pnca-2*), or both transgenes together can strongly rescue the defective MeSa avoidance of the double mutants (Fig. 1B). Though all *nca* transgenes also significantly rescued the locomotion defect, the two *nca* transgenes together appeared to have a stronger effect than either transgene alone (Fig. 1B). Similar to *unc-79(lf)* or *unc-80(lf)* mutants, *nca(lf)* double mutants exhibited normal responses to other attractants and repellents (Fig. S1B).

A gain-of-function (gf) mutation in *nca-1*, *e625*, causes the “coiler” phenotype [23]. To understand how *unc-79* or *unc-80* interacts with *nca-1(e626gf)* in affecting the MeSa avoidance, we generated double mutants. Consistent with previous findings [23], *unc-79(lf)* or *unc-80(lf)* completely suppressed the “coiler” phenotype of the *nca-1(gf)* mutants (unpublished observation). The double mutants also exhibited defective MeSa avoidance (Fig. 1C).

The ER protein NCA localization factor-1 (NLF-1) is required for axonal localization of NCA proteins [35]. To examine whether *nlf-1* is required for the MeSa avoidance, we generated three *nlf-1* deletion lines (Table S2) using a CRISPR/Cas9 method [42]. Taking *nlf-1(mac409)* as a representative lf allele (Fig. 1C), we found that *nlf-1(lf)* itself, or together with *nca-1(lf)*, *nca-2(lf)* or *nca-1(gf)*, did not cause apparently defective MeSa avoidance (Fig. 1C). We did observe that *nlf-1(lf)* suppressed the “coiler” phenotype of *nca-1(gf)* mutants (unpublished observation), consistent with previous findings [35].

To understand how these mutations affect *C. elegans* locomotion, we further compared the body bends of the mutants (Materials and Methods). *unc-79(lf)* and *unc-80(lf)* single mutants, or double mutants carrying either *unc-79(lf)* or *unc-80(lf)*, all exhibited severely defective locomotion (Fig. 1C).

Interestingly, *nca-1(lf)* single mutants exhibited weakly defective locomotion, while *nca-2(lf)* single mutants were moderately defective (Fig. 1C). *nca(lf)* double mutants had severely defective locomotion similar to that of *unc-79(lf)* or *unc-80(lf)* mutants (Fig. 1C).

*nlf-1(lf)* mutants exhibited a moderately defective locomotion, which can be enhanced by *nca-1(lf)* or *nca-2(lf)* (Fig. 1C). However, the defective locomotion of *nlf-1(lf)* mutants appeared to be partially improved by *nca-1(gf)* (Fig. 1C).

### The identification of *unc-80*-expressing neurons

To date, the identities of neurons expressing *unc-80, unc-79* or *nca* remain elusive. To understand the neuronal mechanism underlying the function of *unc-80*, we generated transgenic animals expressing GFP driven by *Punc-80* (Fig. 1A). In these animals, GFP was observed in multiple neurons, ventral nerve cord and vulval muscles (Fig. S3A). A similar expression pattern has been described previously [20, 23].

We next used transgene double-labeling to identify individual neurons that might express *unc-80.* In transgenic animals co-expressing *Punc-80::GFP* (Fig. 2A, left panel) and *Pnmr-1::Cherry* (Fig. 2A, middle panel) (neurons labeled by *Pnmr-1* are listed in Table S3) [43], we observed visible GFP expression in AVA, AVE and AVG neurons (Fig. 2A, right panel) but no obvious expression in AVD and RIM neurons, two classes of neurons reported to be labeled also by *Pnmr-1*. A neuron similar to RIH appeared to be consistently labeled by GFP and mCherry as well (Fig. 2A, right panel). Additional double-labeling using *Punc-80::GFP* (Fig. 2B, 2C and 2D, left panels) with mCherry driven by the *mgl-1* promoter (Table S3) [44, 45] (Fig. 2B, 2C and 2D, middle panels) identified four other classes of GFP-expressing neurons, including RMD and I3 (Fig. 2B, right panel), I4 (Fig. 2C, right panel) and NSM (Fig. 2D, right panel). Except for these neurons, multiple other *Punc-80::GFP*-expressing neurons remain to be identified.

Using a previously described *nlf-1* promoter [35] to drive *mCherry*, we confirmed the neuronal expression pattern of *nlf-1* (Fig. S3B) as previously described [35]. Neuronal double-labeling found that AVA and AVE neurons, among other unidentified neurons, were likely co-labeled by *Punc-80::GFP* and *Pnlf-1::mCherry* (Fig. S4A).

**Figure 2.**
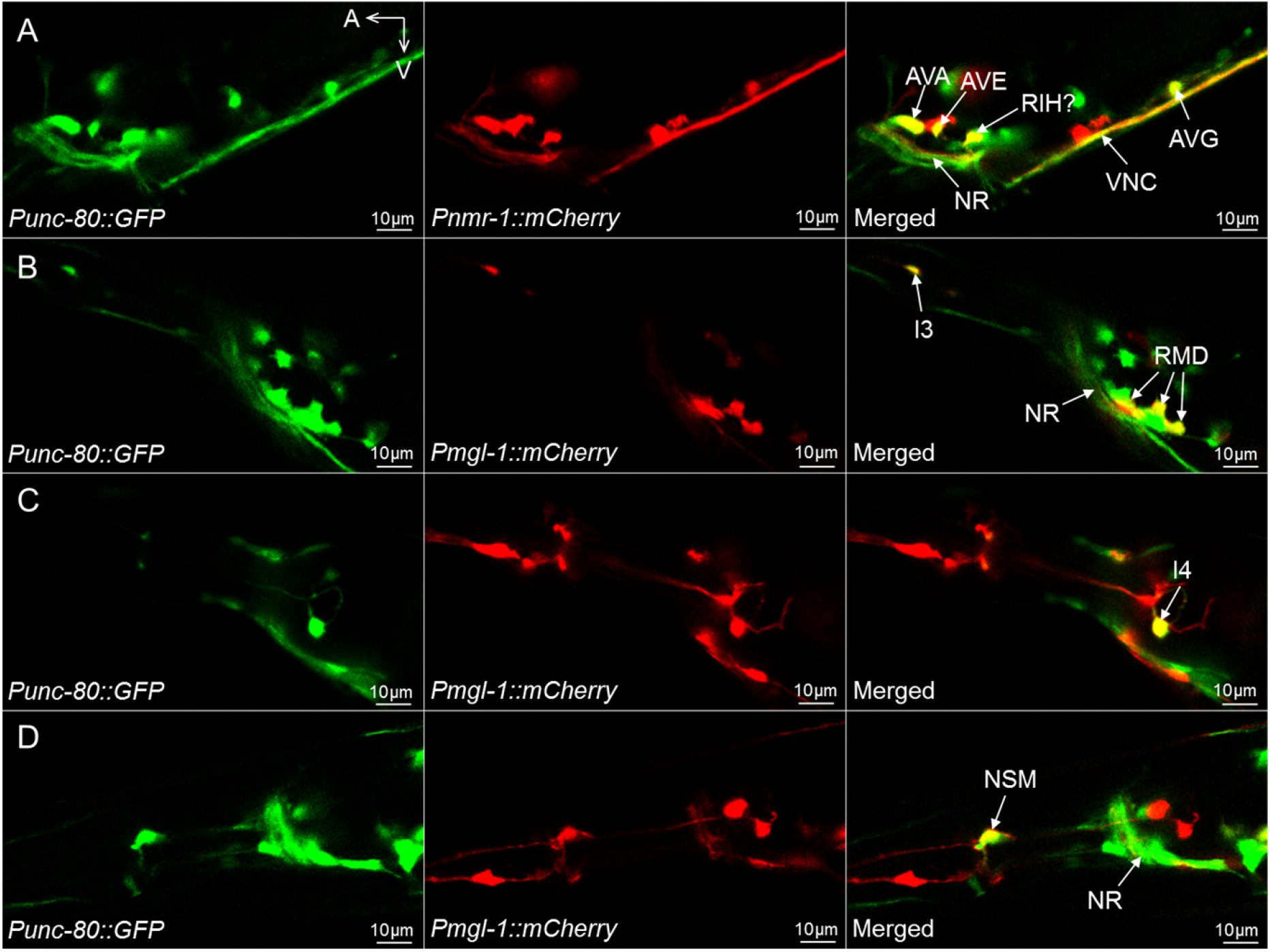
Neuronal double-labeling identified a subset of *unc-80*-expressing neurons. (A) GFP expression driven by the *unc-80* promoter (left panel, *Punc-80*), mCherry driven by the *nmr-1* promoter (middle panel, *Pnmr-1*) and the merged image (right panel) showing GFP expression in AVA, AVE and AVG neurons and potentially in the RIH neuron. (B, C, D) GFP driven by *Punc-80* (left panels), mCherry driven by *Pmgl-1* (middle panels) and the merged images showing GFP expression in RMD and I3 (B, right panel), I4 (C, right panel) and NSM (D, right panel) neurons. VNC: ventral nerve cord. NR: nerve ring. A: anterior. V: ventral. Results were based on three independent transgenic lines. Pictures were taken from a line with the most robust expression of reporters.

### Neuron-specific function of *unc-80*

To understand the function of *unc-80* in different neurons, we performed neuron-specific transgene rescue experiments. An *unc-80* cDNA transgene driven by *Punc-80* can significantly rescue the defective locomotion and MeSa avoidance of *unc-80(lf)* mutants (Fig. 3A, *Punc-80*). Still, the locomotion was partially rescued while the MeSa avoidance was more strongly rescued. The *nlf-1* promoter had similar rescuing effects as the *unc-80* promoter (Fig. 3A, *Pnlf-1*), consistent with the result that *unc-80* and *nlf-1* were co-expressed in multiple neurons (Fig. S4A).

**Figure 3.**
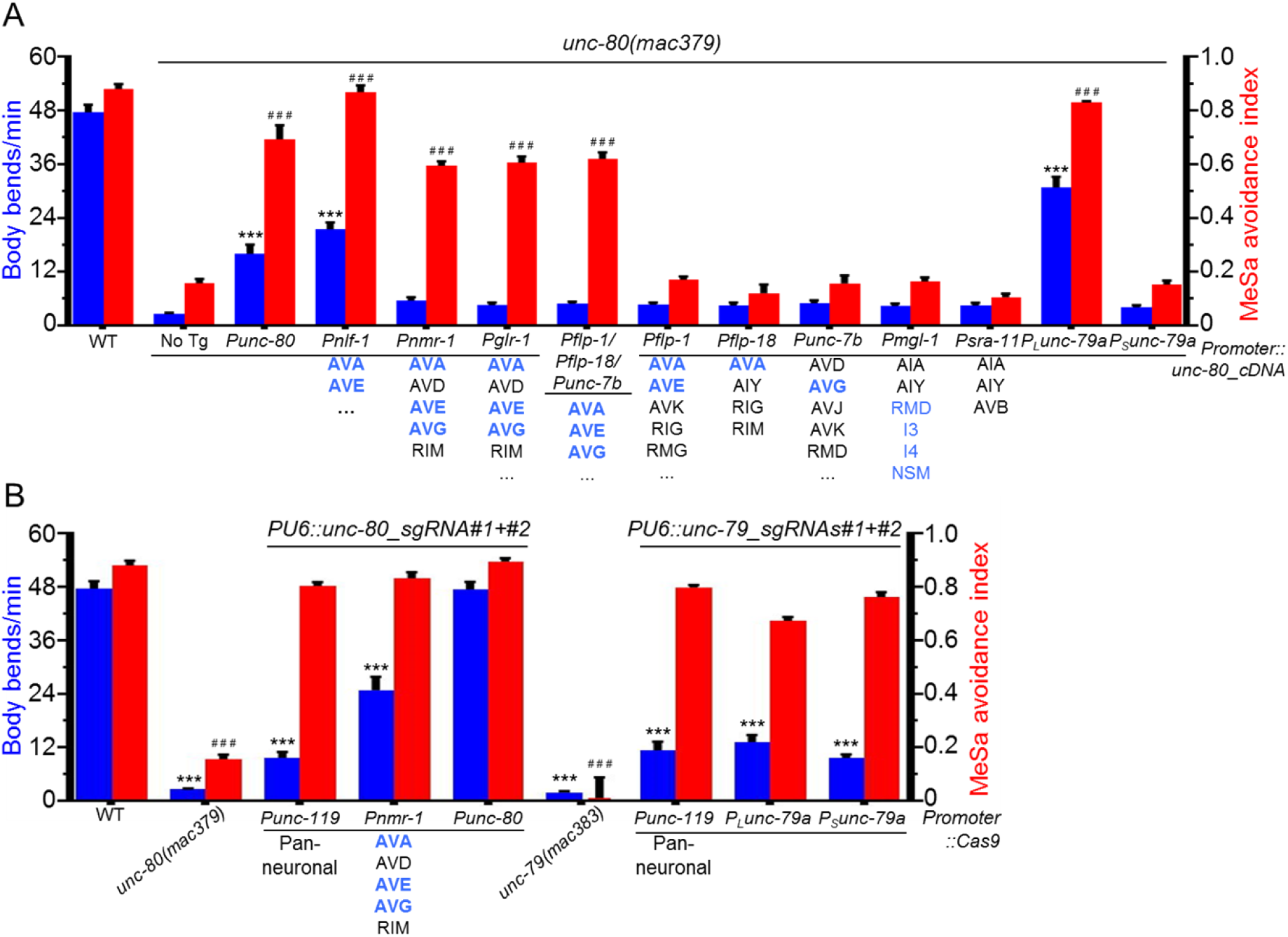
Neuron-specific transgene rescue and neuron-specific knockdown experiments. (A) The locomotion (body bends/min, blue columns, 20 animals per transgenic line, 40 animals in total) and MeSa avoidance indexes (red columns) of *unc-80(lf)* transgenic lines expressing an *unc-80* cDNA transgene driven by different promoters compared to *unc-80(lf)* mutants. *unc-80*-expressing neurons are highlighted in blue. Promoter types and the corresponding neurons are labeled at bottom. Results were based on two independent transgenic lines. (B) The locomotion (body bends/min, blue columns, 20 animals per transgenic line, 40 animals in total) and MeSa avoidance indexes (red columns) of transgenic knockdown lines expressing *Cas9* driven by different promoters and *unc-80*-targeting sgRNAs or *unc-79*-targeting sgRNAs. Results were based on two independent transgenic lines. Comparisons were made with wild type. Statistics: Bonferroni multiple comparison with one-way ANOVA. *** or ^###^, p<0.001. Error bars: standard error of mean. Note: data for wildtype, *unc-79(lf)* and *unc-80(lf)* animals were repeatedly used for comparison purpose.

To examine whether *unc-80* might be effective in a subset of neurons, we used the *nmr-1* promoter (Table S3) to drive the *unc-80* cDNA transgene. The *Pnmr-1::unc-80* transgene significantly rescued the defective MeSa avoidance but not the defective locomotion of *unc-80(lf)* mutants (Fig. 3A, *Pnmr-1*). The *glr-1* promoter (Table S3) [46], which is supposed to drive transgene expression in AVA, AVE and AVG neurons that are also labeled by *Pnmr-1*, similarly rescued the defective MeSa avoidance but not the defective locomotion (Fig. 3A, *Pglr-1*).

To determine whether *unc-80* might be effective in a smaller set of neurons, we tested three other promoters, *flp-1, flp-18* and *unc-7b* (Table S3) [47–49], which label AVA/AVE, AVA, and AVG neurons, respectively. The three *unc-80* transgenes driven by each individual promoter, when injected together, can significantly rescue the defective MeSa avoidance but not the defective locomotion of *unc-80(lf)* mutants (Fig. 3A, *Pflp-1/Pflp-18/Punc-7b*). However, each transgene by itself failed to rescue either defective behavior (Fig. 3A, *Pflp-1*, *Pflp-18* or *Punc-7b*). The *mgl-1* promoter, which drives overlapping expression with *unc-80* in RMD, I3, I4 and NSM neurons (Fig. 2B, 2C and 2D), also failed to rescue either defective behavior (Fig. 3A, *Pmgl-1*). As a negative control, the *sra-11* promoter did not rescue the defective behavior either (Fig. 3A, *Psra-11*), consistent with our finding that its expression did not obviously overlap that of *unc-80* (Fig. S4B).

To understand whether *unc-80* expression in specific neurons is essential for its function, we used a CRISPR/Cas9-based method [50] to examine whether neuron-specific knockdown of *unc-80* would phenocopy the behavioral defects of *unc-80(lf)* mutants.

Transgenic animals expressing two *unc-80*-targeting sgRNAs (Table S4) and *Cas9* driven by the pan-neuronal *unc-119* promoter (Table S3) [51] exhibited obviously defective locomotion (Fig. 3B, *Punc-119*). Limiting the expression of *Cas9* to a subset of interneurons, *e.g.*, AVA, AVE and AVG, using the *nmr-1* promoter also caused defective locomotion (Fig. 3B, *Pnmr-1*). However, *Punc-80::Cas9* failed to cause obviously defective locomotion (Fig. 3B, *Punc-80*). We postulate that the expression of *Punc-80*::*Cas9* might not be strong enough in disrupting the *unc-80* locus in these animals.

Surprisingly, different from the locomotion, the MeSa avoidance response was not affected in any of the knockdown lines (Fig. 3B, red columns).

### *unc-79* was largely co-expressed with *unc-80*

Though it is well accepted that UNC-79 and UNC-80 function together, it remains unclear whether *unc-79* and *unc-80* are co-expressed in the same neurons. To answer this question, we generated transgenic lines expressing GFP driven by the *P_L_unc-79a* or *P_S_unc-79a* promoter (Fig. 1A). GFP driven by *P_L_unc-79a* was expressed in multiple head and tail neurons, ventral nerve cord and vulval muscles (Fig. S3C), a pattern similar to that of *unc-80* (Fig. S3A). However, GFP driven by *P_S_unc-79a* was only detected in several pairs of head neurons (Fig. S3D). *P_S_unc-79::GFP* was also observed in anterior and posterior portions of the intestine (Fig. S3D), which we consider to be non-specific as *P_L_unc-79::GFP* was not detected there (Fig. S3C).

To investigate whether *unc-79* and *unc-80* were co-expressed, we generated transgenic animals co-expressing *P_L_unc-79a::GFP* and *Punc-80::mCherry*. In these animals, GFP and mCherry co-labeled a large number of head neurons (Fig. 4A), the vulval muscles (Fig. 4B) and a few tail neurons (Fig. 4C). At the same time, one or more head neurons (Fig. 4A, right panel, arrow heads), motor neuron (Fig. 4B, right panel, arrow head) or tail neuron (Fig. 4C, right panel, arrow head) appeared to be labeled by only GFP or mCherry but not both. Consistent with the largely overlapping expression pattern, a *P_L_unc-79::unc-80* cDNA transgene significantly rescued the defective locomotion and MeSa avoidance of *unc-80(lf)* mutants (Fig. 3A, *P_L_unc-79*).

**Figure 4.**
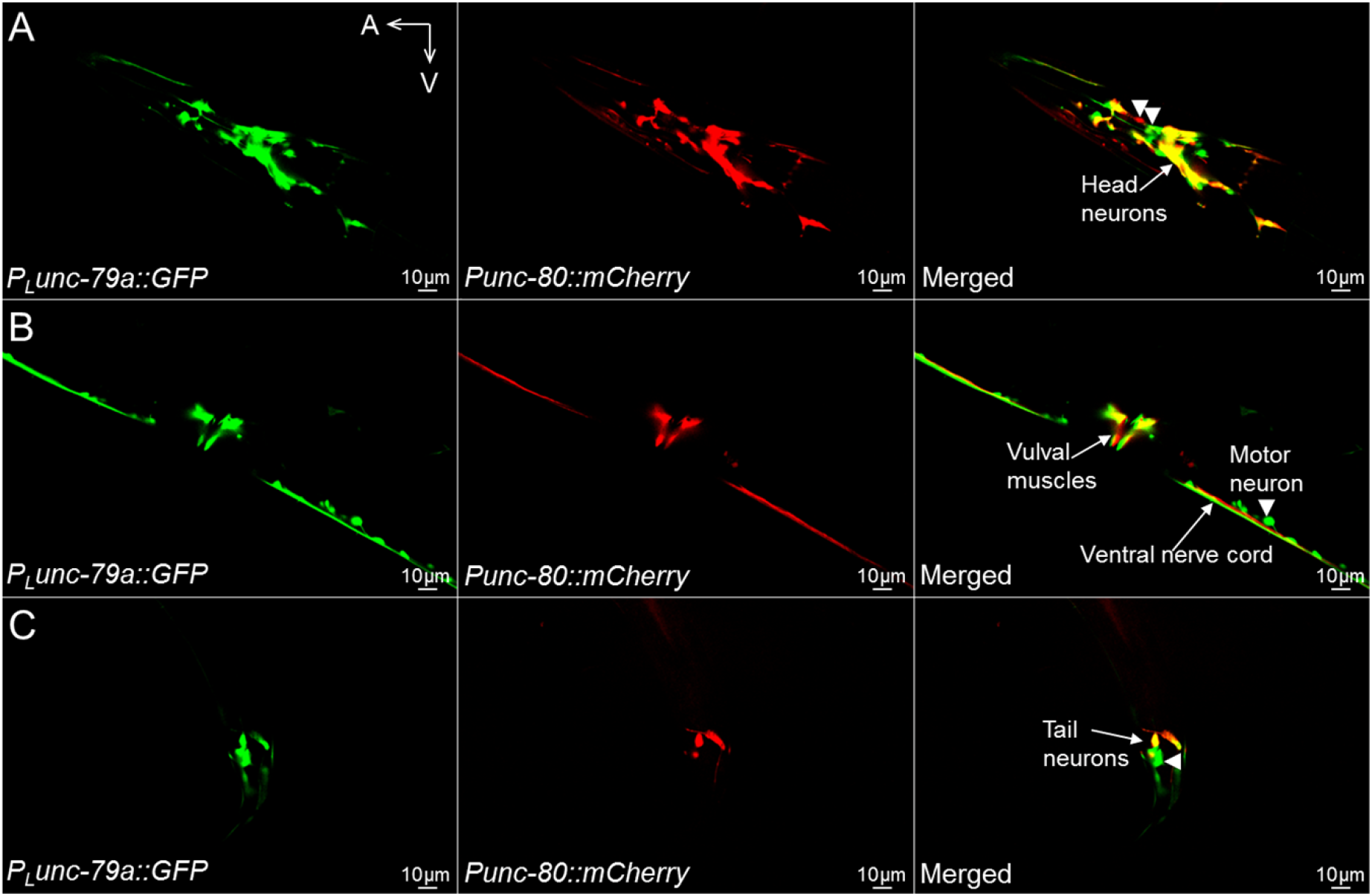
Neuronal double-labeling found that *unc-79* expression largely overlaps that of *unc-80*. GFP expression driven by the long *unc-79a* promoter (left panel, *P_L_unc-79a*), mCherry driven by the *unc-80* promoter (middle panel, *Punc-80*) and the merged image (right panel) showing multiple head neurons co-expressing GFP and mCherry. Same as (A), showing vulval muscles co-labeled by GFP and mCherry. (C) Same as (A, B), showing a few tail neurons co-labeled. Arrow heads point to neurons that appear to express only one fluorescent reporter. A: anterior. V: ventral. Results were based on three independent transgenic lines. Pictures were taken from a line with the most robust expression of reporters.

The limited expression pattern of the *P_S_unc-79a* promoter (Fig. S3D) and its partial rescuing power (Fig. 1B) suggest that it might define a subset of *unc-79*-expressing neurons. We combined DiI tracing and neuronal double-labeling to identify the neurons labeled by this promoter.

In *P_S_unc-79::GFP* transgenic animals, we found that GFP co-labeled ASH and ASJ that were also stained with DiI (Fig. 5A). The expression of *P_S_unc-79::GFP* (Fig. 5B and 5C, left panels) in ASJ (Fig. 5B, right panel) and ASH (Fig. 5C, right panel) neurons was verified by co-labeling with *Pssu-1::mCherry* (Table S3) [52] (Fig. 5B, middle panel) and *Psra-6::mCherry* (Table S3) [53] (Fig. 5C, middle panel), respectively. *P_S_unc-79::GFP* (Fig. 5D, left panel) also appeared to label the RIA neurons (Fig. 5D, right panel), which were identified by co-labeling with *Pglr-3::mCherry* (Table S3) [46] (Fig. 5D, middle panel). The fourth pair of neurons labeled by *P_S_unc-79::GFP* was similar to the closely positioned motor neurons RMF or RMH (Fig. S4C), which remains to be clarified. Unlike the long *unc-79* promoter (Fig. 3A, *P_L_unc-79a*), a *P_S_unc-79a::unc-80* transgene failed to rescue the defective locomotion or MeSa avoidance of *unc-80(lf)* mutants (Fig. 3A, *P_s_unc-79*), consistent with the finding that *P_S_unc-79a::mCherry* and *Punc-80::GFP* did not appear to co-label neurons (Fig. S4D).

**Figure 5.**
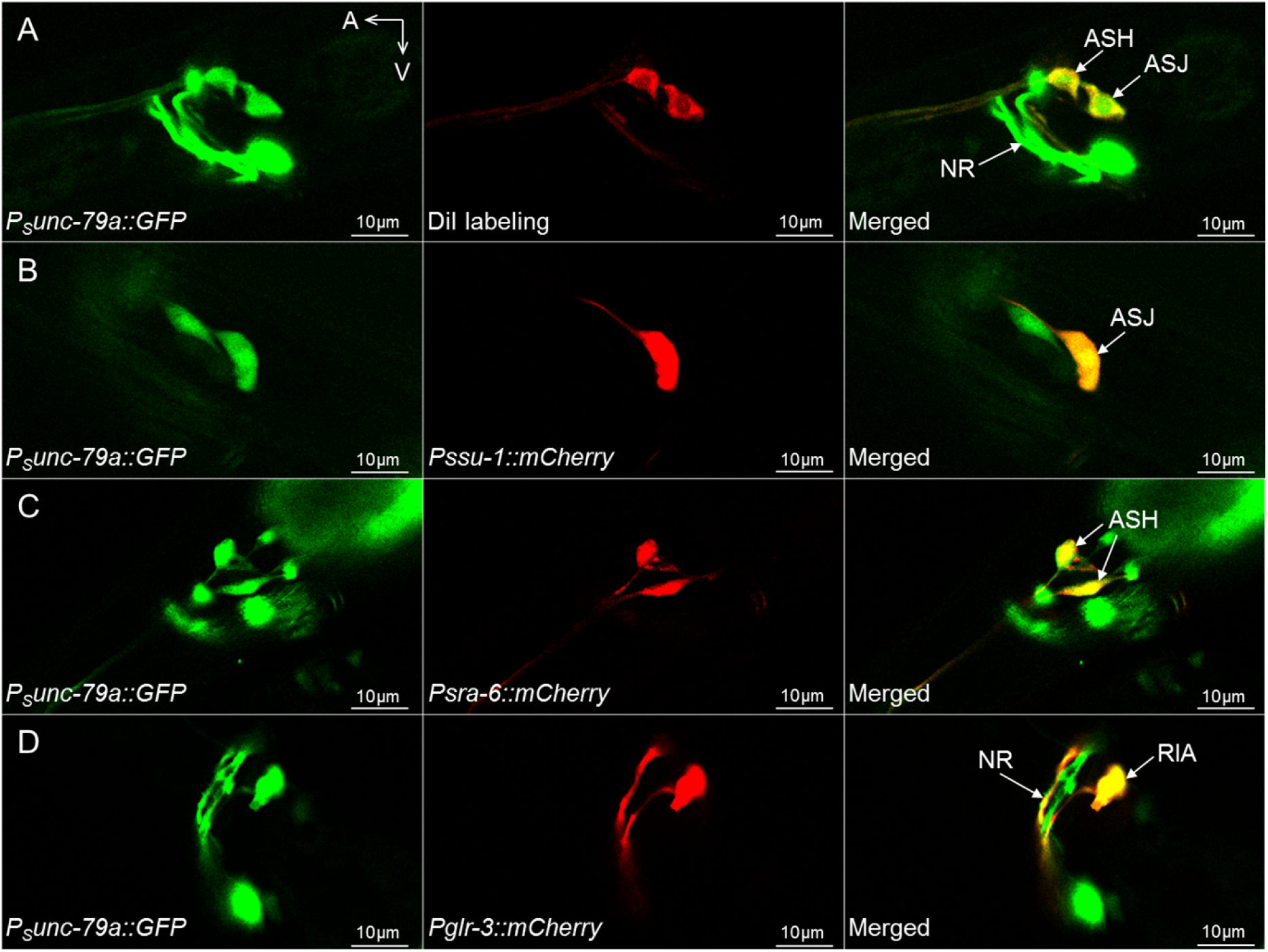
DiI tracing and neuronal double-labeling identified neurons potentially labeled by the short *unc-79a* promoter. (A) GFP expression driven by the short *unc-79a* promoter (left panel, *P_S_unc-79a*), DiI labeling of sensory neurons (middle panel) and the merged image (right panel) showing GFP expression in ASJ and ASH neurons. (B, C, D) GFP driven by *P_S_unc-79a* (B, C, D, left panels), mCherry driven by *Pssu-1* (B, middle panel), *Psra-6* (C, middle panel) or *Pglr-3* (D, middle panel) and the merged images showing GFP expression in ASJ neurons (B, right panel), ASH neurons (C, right panel) and RIA neurons (D, right panel). NR: nerve ring. A: anterior. V: ventral. Results were based on three independent transgenic lines. Pictures were taken from a line with the most robust expression of reporters.

We next used the CRISPR/Cas9 knockdown to investigate the function of *unc-79* in neurons. Transgenic animals expressing two *unc-79-*targeting sgRNAs (Table S4) and *Punc-119::Cas9* exhibited significantly defective locomotion (Fig. 3B, *Punc-119*). *Cas9* driven by the long or short *unc-79* promoter also caused defective locomotion (Fig. 3B, *P_L_unc-79a and P_S_unc-79a*). However, none of the *unc-79*-knockdown lines exhibited obviously defective MeSa avoidance (Fig. 3B, red columns), which is similar to *unc-80-*knockdown animals.

## Discussion

In this study, we provide genetic evidence that *unc-79*, *unc-80* and the *nca* genes are specifically required for *C. elegans* avoidance response to the plant hormone methyl salicylate. *unc-79* and *unc-80* are expressed in largely overlapping neurons to affect animal behavior. We suggest that interneurons AVA, AVE and AVG might play key roles in mediating the functions of *unc-80*.

Studies in mammals suggest that UNC79, UNC80 and NALCN can interact biochemically. In mouse brain or human HEK293 fibroblast cells, NALCN and UNC80 can form a complex [26]. This complex also interacts with UNC79 [27]. In *C. elegans*, UNC-80 and UNC-79 are required for axonal localization of the NCA channel [20, 23]. UNC79 also affects the expression levels of NALCN in *C. elegans* and *Drosophila* [19].

We found that *C. elegans nca*, *unc-79* and *unc-80* lf mutants exhibited similar MeSa-specific avoidance defects, while these mutants had similar wild type-like responses to other repellents or attractants (Fig. 1C and S1B). *unc-79* and *unc-80* were co-expressed in multiple neurons (Fig. 4). Consistently, an *unc-80* cDNA transgene driven by the long *unc-79* promoter can significantly rescue the defects of *unc-80(lf)* mutants (Fig. 3A). Therefore, our findings provide novel genetic and neuronal evidence that NALCN, UNC80 and UNC79 function together to regulate neuronal activities.

The transgenic rescue and knockdown experiments suggest that *unc-80* expression in interneurons AVA, AVE and AVG might be sufficient and necessary for the MeSa avoidance (Fig. 3), though the effects of other *unc-80-*expressing neurons remain to be determined. The key role of AVA neurons in mediating *unc-80* functions was also suggested by Gao et al, who found that NCA can activate the AVA neurons to potentiate persistent motor circuit activity in *C. elegans* [21].

Multiple studies found that AVA neurons play a central role in a variety of *C. elegans* behaviors, *e.g.*, touch-induced movement [54, 55], reversal locomotion [56], mechanosensory habituation [57], variability in reversal response to odor stimuli [58] and repetitive reversals caused by glutamate spillover [59]. That AVA excitation is required for reversal locomotion provides an interpretation for the actions of *nca*, *unc-80* and *unc-79* in the MeSa avoidance, for it is plausible that loss-of-function in these genes might cause hyperpolarization of AVA neurons, which would lead to AVA inactivation and subsequent lack of reversals in response to MeSa.

We previously found that multiple neuronal genes might affect the MeSa avoidance response [36]. Neuron-specific rescue experiments suggest that the *npr-1*-expressing inter/motor neurons RMG and the *npr-2*-expressing interneurons AIZ might be involved [36]. Among sensory neurons, AWB appears to play a key role in detecting MeSa, while AWC might play a minor role [36]. Together with the current study, our results suggest the involvement of the AWC, AIZ and AVA neurons in mediating the MeSa avoidance response. This model is consistent with the placement of these neurons in a core circuit that mediates *C. elegans* chemotaxis [58]. In the presence of MeSa, the AVE and AVG interneurons might also communicate with this core circuit to elicit the avoidance response.

The *nca*, *unc-80* and *unc-79* lf mutants exhibited MeSa-specific defective avoidance but had wild type-like responses to other repellents or attractants (Fig. S1B). This difference implies that *C. elegans* might express a MeSa-specific receptor. Previously, the human TRPV1 channel was shown to be inhibited by MeSa [60]. However, the wild type-like response to MeSa by two TRPV channel mutants (*osm-9* and *ocr-2*) [36] and the finding that all five TRPV channel expression was not detected in AWB neurons [61, 62] do not support a TRPV channel as the MeSa receptor in *C. elegans*. Alternatively, a GPCR expressed in antenna sensory neurons of the tortricid moth was found to exhibit high sensitivity to MeSa in the insect sf9 cells [63]. This result is consistent with the findings that GPCRs can regulate NALCN activities in mammals [9,26–28,64], NCA are downstream targets of neuronal G-protein signals in *C. elegans* [29, 30] and GPCR signals are involved in *C. elegans* avoidance response to MeSa [36]. We also suspect that MeSa might affect *C. elegans* behavior by directly activating the NALCN channel, *e.g.*, by activating NALCN expressed on sensory neurons or interneurons that mediate the reversal locomotion. Therefore, future studies are warranted for understanding the underlying molecular mechanisms.

It is interesting that *nca* single mutants exhibited partially defective locomotion but largely normal MeSa avoidance (Fig. 1C), so did the *nlf-1(lf)* mutants (Fig. 1C). Similarly, Neuron-specific knockdown of *unc-79* or *unc-80* significantly affected the locomotion but not the MeSa avoidance (Fig. 3B). Furthermore, *unc-80* transgenes driven by *nmr-1*, *glr-1* or *flp-1/flp-18/unc-7b* promoters significantly improved the MeSa avoidance but not the locomotion of *unc-80(lf)* mutants (Fig. 3A). Though puzzling, we may find that a common feature of these strains is their similarity to partial loss-of-function mutants, because these animals express *nca*, *unc-79* or *unc-80* at levels between that of null mutants and wild type. Hence, compared to the MeSa avoidance, the locomotion behavior appears to be more dependent on the expression levels of these genes, suggesting that it provides a more sensitive assay for analyzing the functions of *nca*-related genes.

The short *unc-79a* promoter (*P_s_unc-79a*) exhibited a more limited expression pattern, similar to what was mentioned previously [19]. *P_s_unc-79a* was capable of rescuing the defective MeSa avoidance of *unc-79(lf)* mutants (Fig. 1B). However, when driving an *unc-80* cDNA transgene, this promoter failed to obviously rescue *unc-80(lf)* mutants (Fig. 3A). Such a difference might reflect the non-overlapping expression of *P_s_unc-79a* with *Punc-80* (Fig. S4D). Alternatively, it might be caused by the structure of the transgenes. To rescue *unc-79(lf)* mutants, *P_s_unc-79a* was driving an *unc-79a* gDNA transgene (Fig. 1A), while to rescue *unc-80(lf)* mutants, the promoter was driving an *unc-80* cDNA transgene (Fig. 3A). In the formal situation, the *unc-79* gDNA transgene might carry transcriptional regulatory elements that can compensate the limited expression of *P_s_unc-79a*. However, in the latter situation, the *unc-80* cDNA transgene would not carry compensating regulatory elements and would be insufficient for the rescue. Nevertheless, more experiments are needed to determine whether the expression pattern of the short *unc-79* promoter provides a neuronal mechanism underlying certain *unc-79* functions.

Both UNC-79 and UNC-80 contain a string of armadillo-repeat domains with unknown functions (Fig. S2). Armadillo repeats are ∼42 amino acid motifs that have been found in signaling, cytoskeleton and structural proteins to mediate a broad array of protein-protein interactions [65]. The armadillo repeats in UNC-79 and UNC-80 might mediate interactions with unidentified proteins that could add extra layers of regulation on NALCN.

In short, we found that *unc-79*, *unc-80* and *nca* genes are required specifically for *C. elegans* avoidance response to the plant hormone MeSa. *unc-79* and *unc-80* are expressed and function in largely overlapping neurons. The AVA, AVE and AVG interneurons appear to be necessary and sufficient for *unc-80* regulation of the MeSa avoidance. Our findings uncovered novel functions of the NALCN channel complex and provide new molecular and neuronal mechanisms for understanding the regulation of the NALCN channel.

## Materials and Methods

### Strains

See supplementary Materials and Methods.

### MeSa avoidance assay

*C. elegans* MeSa avoidance assay was performed as previously described [36]. 30 to 200 animals were examined in each experiment.

### Locomotion assay

Synchronized L4 animals were picked into new plates seeded with OP50 bacteria one night before the assay. Body bends were measured by touching an animal on the tail with a worm pick to help initiate locomotion, followed by counting the number of body bends (one head turn) for 1 min.

### Genetic screen for and identification of mutants with defective avoidance response to MeSa

Synchronized L4 wild-type animals (P_0_) were mutagenized with EMS (ethyl methane sulfonate) as described [66]. F_1_ progeny were allowed to grow to young adults and bleached to generate synchronized F_2_ progeny for the MeSa avoidance assay. Adult F_2_ animals that failed to avoid MeSa were collected from each individual plate and bleached to generate synchronized adult F_3_ progeny for a new round of MeSa assay. After six rounds of selection, an individual F_7_ progeny that failed to avoid MeSa was picked from each plate, allowed to propagate and retested in the MeSa avoidance assay. From ∼60,000 F_1_ animals, we isolated 15 independent mutants.

Genetic complementation tests using the response of F_1_ males to MeSa as readout identified three distinct groups, with group one containing 11 mutants (*mac379, mac380, mac381, mac382, mac384, mac385, mac386, mac388, mac390, mac391, mac394*), group two containing three mutants (*mac383, mac389, mac393*) and group three containing one mutant (*mac387*) (Table S1). We selected *mac379*, *mac382* and *mac388* from group one, and *mac383* from group two for genomic sequencing as described [67]. Sequencing results indicated that *mac379*, *mac382* and *mac388* carried distinct nonsense mutations in the *unc-80* gene and *mac383* carried a nonsense mutation in *unc-79*. The *mac387* mutation in the third complementation group was not further analyzed, as it appeared to affect general chemotaxis of *C. elegans* (Fig. S1B).

### Molecular biology

See supplementary Materials and Methods. PCR primers are listed in Table S5.

### Transgene experiments

Germline transgene experiments were performed as described [68].

For genomic transgene rescue experiments, transgenic mixtures contained 10∼20 ng/μl genomic PCR fragment and 20 ng/μl pPD95_86 (*Pmyo-3::GFP*) as co-injection marker.

For transcriptional reporter experiments, transgene solutions containing 20∼50 ng/μl reporter construct were injected to wild type.

For neuron-specific knockdown experiments, we used a previously described method with minor modifications [50]. The transgene mixtures contained 20∼50 ng/μl *Promoter::Cas9::NLS::3’UTR*, 25 ng/μl *PU6::unc-79_sgRNA#1* and *#2* each *(or PU6::unc-80_sgRNA#1* and *#2* each), and 20 ng/μl pPD95_86 (*Pmyo-3::GFP*) or 2.5 ng/μl pCFJ90 (*Pmyo-2::mCherry*) as co-injection marker.

For neuron-specific rescue experiments, transgenic mixtures contained 10∼20 ng/μl *Promoter::unc-80_cDNA* and 20 ng/μl pPD95_86 (*Pmyo-3::GFP*) or 2.5 ng/μl pCFJ90 (*Pmyo-2::mCherry*) as co-injection marker.

### Identification of *unc-79-* and *unc-80-*expressing neurons

We used DiI-labeled sensory neurons [69] as landmarks to facilitate the identification of *unc-79*-expressing neurons. Images of fluorescence-positive neurons in transgenic animals expressing GFP and/or mCherry reporters were captured using a 63X DIC/fluorescent Leica TCS SP5 II laser confocal microscope and neuronal identities were inferred by overlapping fluorescence signals and by comparing to the anatomical and morphological characteristics of neurons described in Wormatlas (www.wormatlas.org).

### Statistical analysis

*P* values were determined by Paired two-tailed Student’s *t*-test or Bonferroni’s multiple comparison using GraphPad Prism 7.0 software.

## Acknowledgments

We thank members of the Ma laboratory for suggestions. Some strains were provided by the CGC, which is funded by NIH Office of Research Infrastructure Programs (P40 OD010440).

## Author contributions

CZ and LM designed the experiments. CZ performed the genetic, molecular and neuronal analyses with the assistance of JL, XH, QZ, YH and XW. CZ and LM wrote the manuscript. LM managed the project.

## Data availability

Relevant data are within the manuscript and its Supporting information files.

## Supplementary Figures and Tables

**Figure S1.**
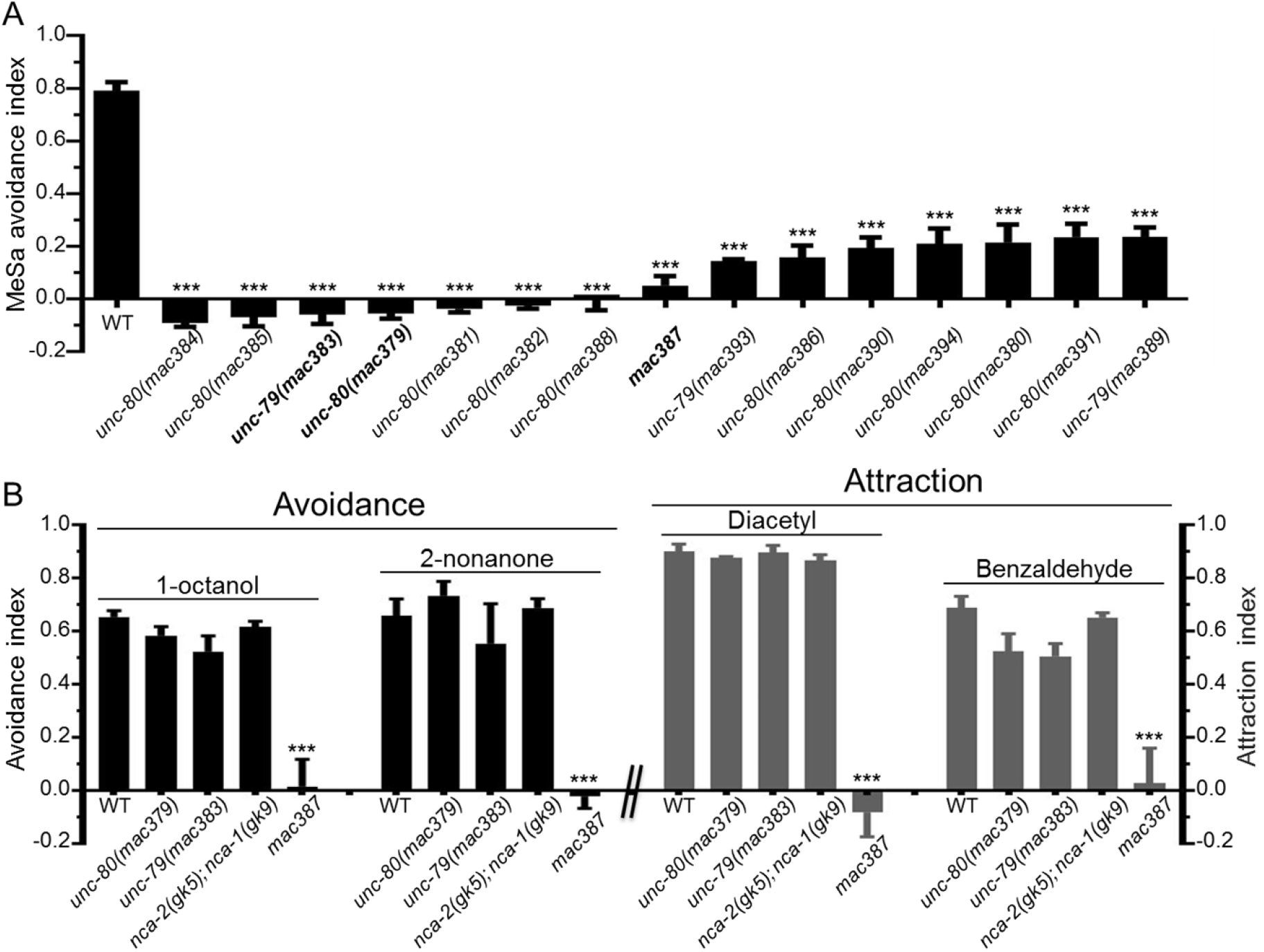
Chemotaxis of mutants with defective MeSa avoidance responses. (A) MeSa avoidance indexes of mutants isolated in the MeSa avoidance mutant screening. (B) Responses of representative mutants to volatile repellents and attractants, including repellents 1-octanol and 2-nonanone and attractants diacetyl and benzaldehyde. Each dataset was based on three biological replicates. Statistics: Bonferroni multiple comparison with one-way ANOVA. ***, p<0.001. Error bars: standard error of mean.

**Figure S2.**
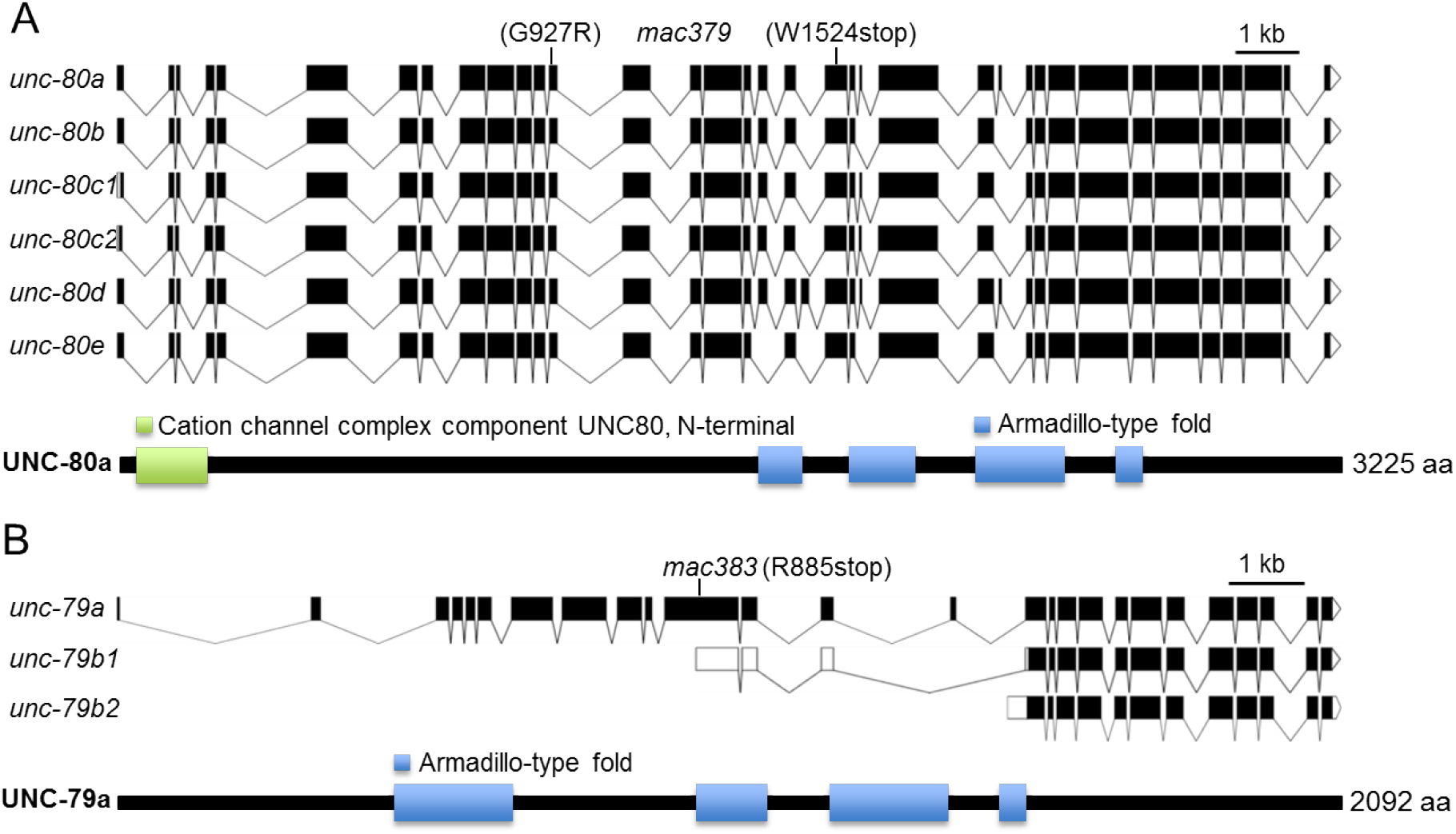
Exon-intron structures of *unc-80* and *unc-79* and protein domains of UNC-80 and UNC-79. (A) *unc-80* gene structure and transcript isoforms, UNC-80 protein domains and the position of the *mac379* lf mutation. (B) *unc-79* gene structure and transcript isoforms, UNC-79 protein domains and the position of the *mac383* lf mutation. Protein domains were identified based on sequence searches at www.ebi.ac.uk/interpro.

**Figure S3.**
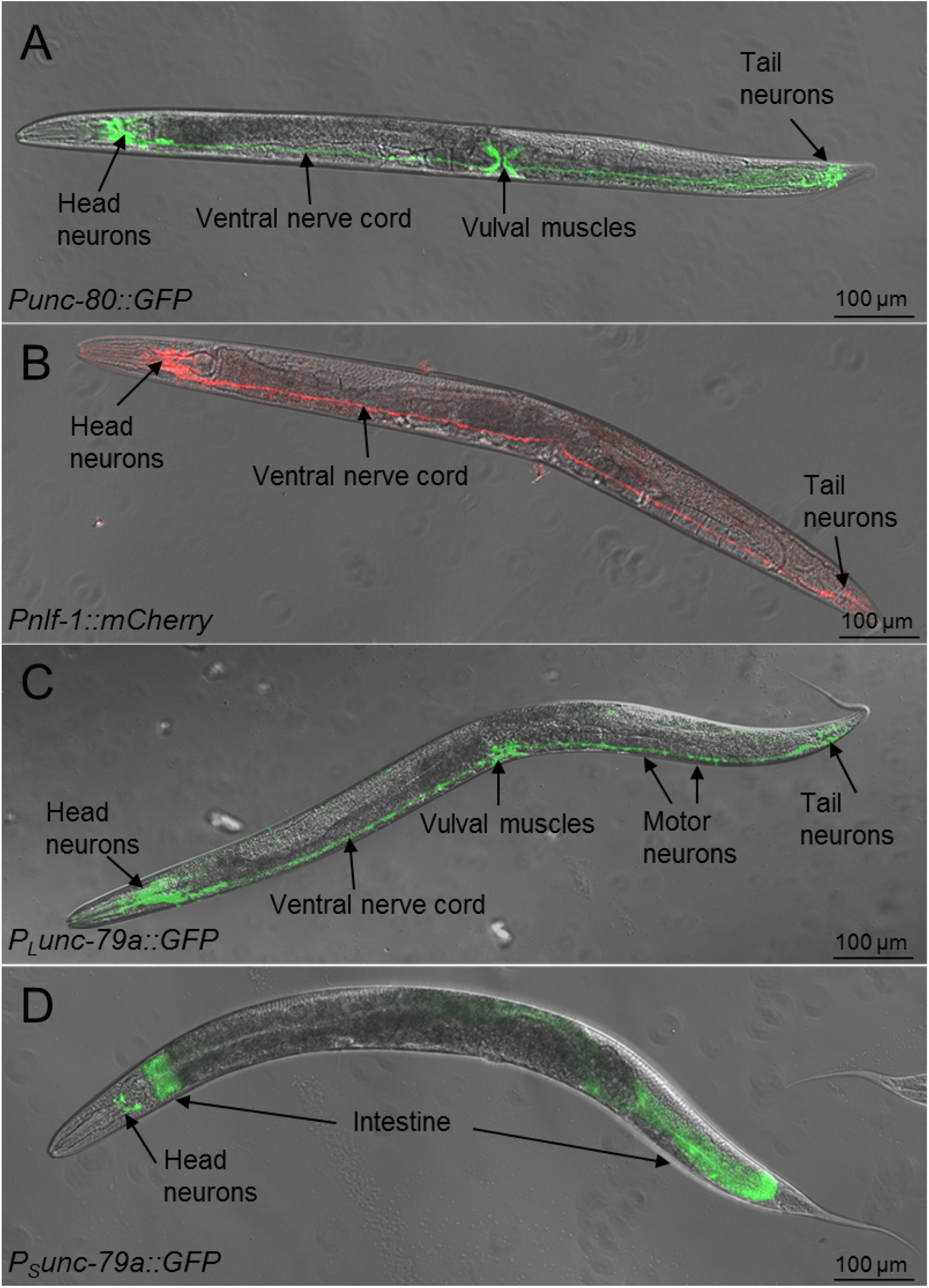
Representative pictures of transgenic animals expressing fluorescent reporters driven by *unc-80, nlf-1* or *unc-79* promoters. (A) A *Punc-80::GFP* transgenic adult showing GFP expression in head neurons, tail neurons, ventral nerve cord and vulval muscles (B) A *Pnlf-1::mCherry* transgenic adult showing mCherry expression in head neurons and ventral nerve cord. (C) A *P_L_unc-79a::GFP* transgenic adult showing GFP expression in head neurons, motor neurons, tail neurons, ventral nerve cord and vulval muscles. (D) A *P_S_unc-79a::GFP* transgenic adult showing GFP expression in a few head neurons and the anterior and posterior portions of the intestine.

**Figure S4.**
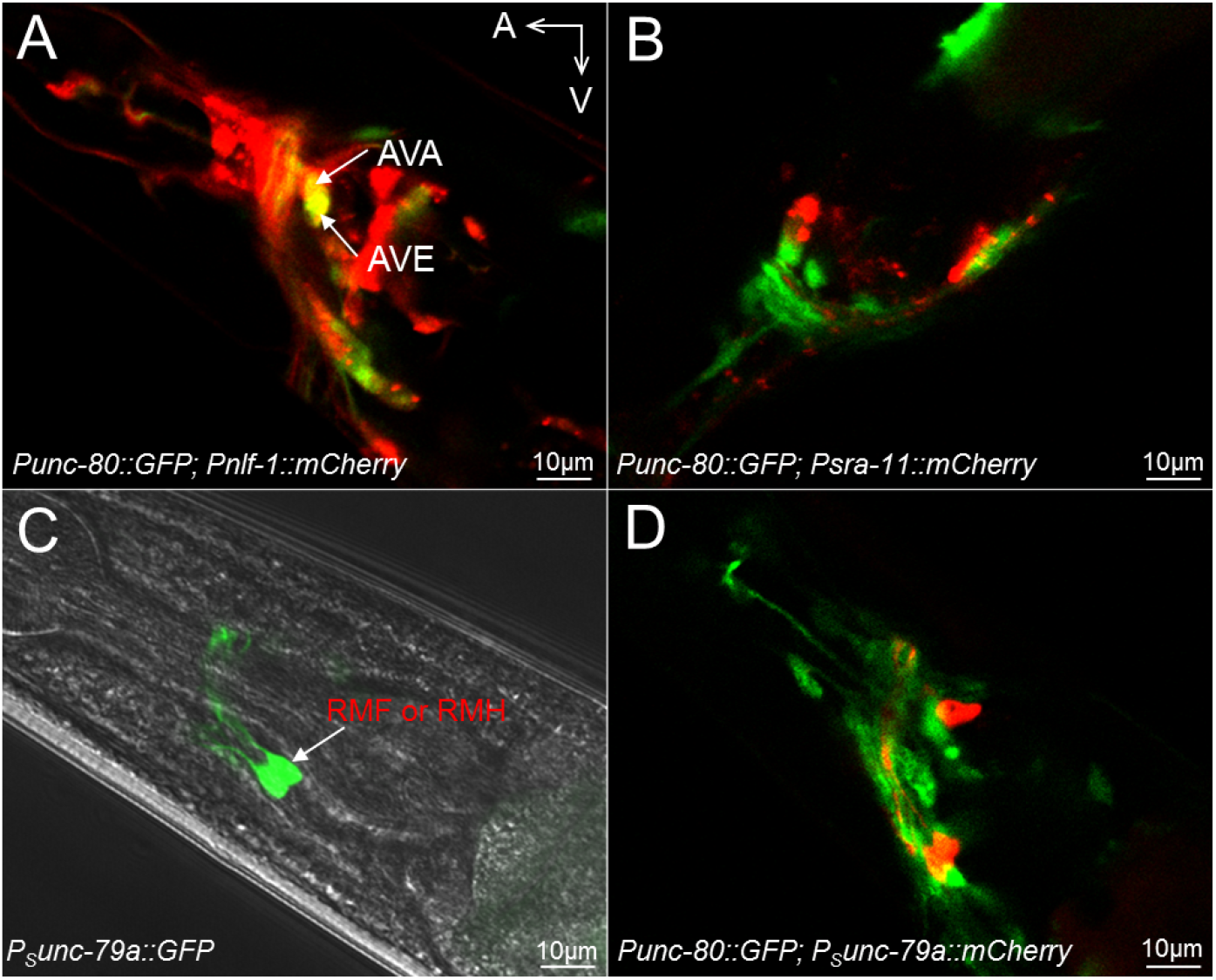
Transgenic animals with fluorescent protein expression driven by different promoters. (A) A confocal image of a transgenic adult expressing *Punc-80::GFP* and *Pnlf-1::mCherry* showing AVA and AVE neurons co-labeled by GFP and mCherry. (B) A confocal image of a transgenic adult expressing *Punc-80::GFP* and *Psra-11::mCherry* showing no neurons obviously co-labeled by GFP and mCherry. (C) RMF- or RMH-like neurons expressing *P_S_unc-79a::GFP* in a phase contrast background. (D) A confocal image of a transgenic adult expressing *Punc-80::GFP* and *P_S_unc-79a::mCherry* showing no neurons obviously co-labeled by GFP and mCherry. A: anterior. V: ventral.

**Table S1.** List of *unc-80* and *unc-79* mutations isolated in the MeSa avoidance mutant screening.

| Group | Gene | Allele | Exon/Intron | Mutation | Effect |
| --- | --- | --- | --- | --- | --- |
| 1 | *unc-80a* | ***mac379*** | Exon 13 | GGA→AGA | G927R |
|  |  |  | Exon 20 | TGG→TGA | W1524stop |
|  | *unc-80a* | *mac380* | Exon 4 | CCG→TAG | P109stop |
|  | *unc-80a* | *mac381* | Exon 31 | CAG→TAG | Q2740stop |
|  | *unc-80a* | ***mac382*** | Exon 6 | TGG→TGA | W220stop |
|  | *unc-80a* | *mac384* | Intron 33 | GC→AC | 5' splice site |
|  | *unc-80a* | *mac385* | Exon 27 | CAG→TAG | Q2043stop |
|  | *unc-80a* | *mac386* | Exon 33 | CGA→TGA | R2940stop |
|  | *unc-80a* | ***mac388*** | Exon 24 | TGG→TAG | W1967stop |
|  | *unc-80a* | *mac390* | Exon 16 | TGG→TGA | W1292stop |
|  | *unc-80a* | *mac391* | Exon 23 | CGA→TGA | R1886stop |
|  | *unc-80a* | *mac394* | Exon 31 | TGG→TGA | W2709stop |
| 2 | *unc-79a* | ***mac383*** | Exon 11 | CGA→TGA | R885stop |
|  | *unc-79a* | *mac389* | Exon 15 | 56 bp deletion | Frameshift |
|  | *unc-79a* | *mac393* | Intron 2 to Exon 4 | 508 bp deletion | Frameshift |
| 3 | ND | ***mac387*** | ND | ND | ND |

Representative alleles used for chemotaxis experiments are listed in bold. Positions of mutations, nucleotide changes and effects on amino acids are indicated. ND: not determined. *unc-80(mac379)* and *unc-79(mac383)* were used as reference If alleles in this study.

**Table S2.** List *nlf-1* deletion mutations generated using the CRISPR/Cas9 method.

| *nlf-1* allele | Mutation | Effect | Isolation method |
| --- | --- | --- | --- |
| *mac408* | 30 bp deletion | 29 nt deletion in exon 4 and intron 4 3' SS | CRISPR/Cas9 |
| *mac409* | 33 bp deletion | 27 nt deletion in exon 4 and intron 4 3' SS | CRISPR/Cas9 |
| *mac410* | 36 bp deletion | 22 nt deletion in exon 4 and intron 4 3' SS | CRISPR/Cas9 |

*mac409* was used as reference *lf* allele in this study. *nlf-1* sgRNA target sequence is shown in Table S4.

**Table S3.** List of neuron-specific promoters and the classes of neurons labeled by each promoter.

| Promoter | Expression |
| --- | --- |
| *nmr-1* | **AVA**, AVD, **AVE**, **AVG**, PVC, RIM, **RIH?** |
| *mgl-1* | AIA, AIY, NSM, RMD, **I3**, **I4** |
| *glr-1* | **AVA**, AVB, AVD, **AVE**, PVC, AIB, RMD, RIM, SMD, **AVG**, PVQ, URY |
| *flp-1* | **AVA**, **AVE**, AVK, RIG, RMG, AIA, AIY, M5 |
| *flp-18* | **AVA**, AIY, RIG, RIM |
| *unc-7b* | AVD, **AVG**, AVJ, AVK, DVA, DVC, IL1, PVP, RIC, RIF, RMD, SIB, SMD, URA |
| *sra-11* | AIA, AIY, AVB, PVT, RIF, RIG |
| *unc-119* | Pan-neuronal |
| *ssu-1* | ASJ |
| *sra-6* | ASH, ASI, PVQ |
| *glr-3* | RIA |

Promoter expression patterns are based on annotations in www.wormbase.org. For some promoters, not all annotated neurons are shown.

**Table S4.**
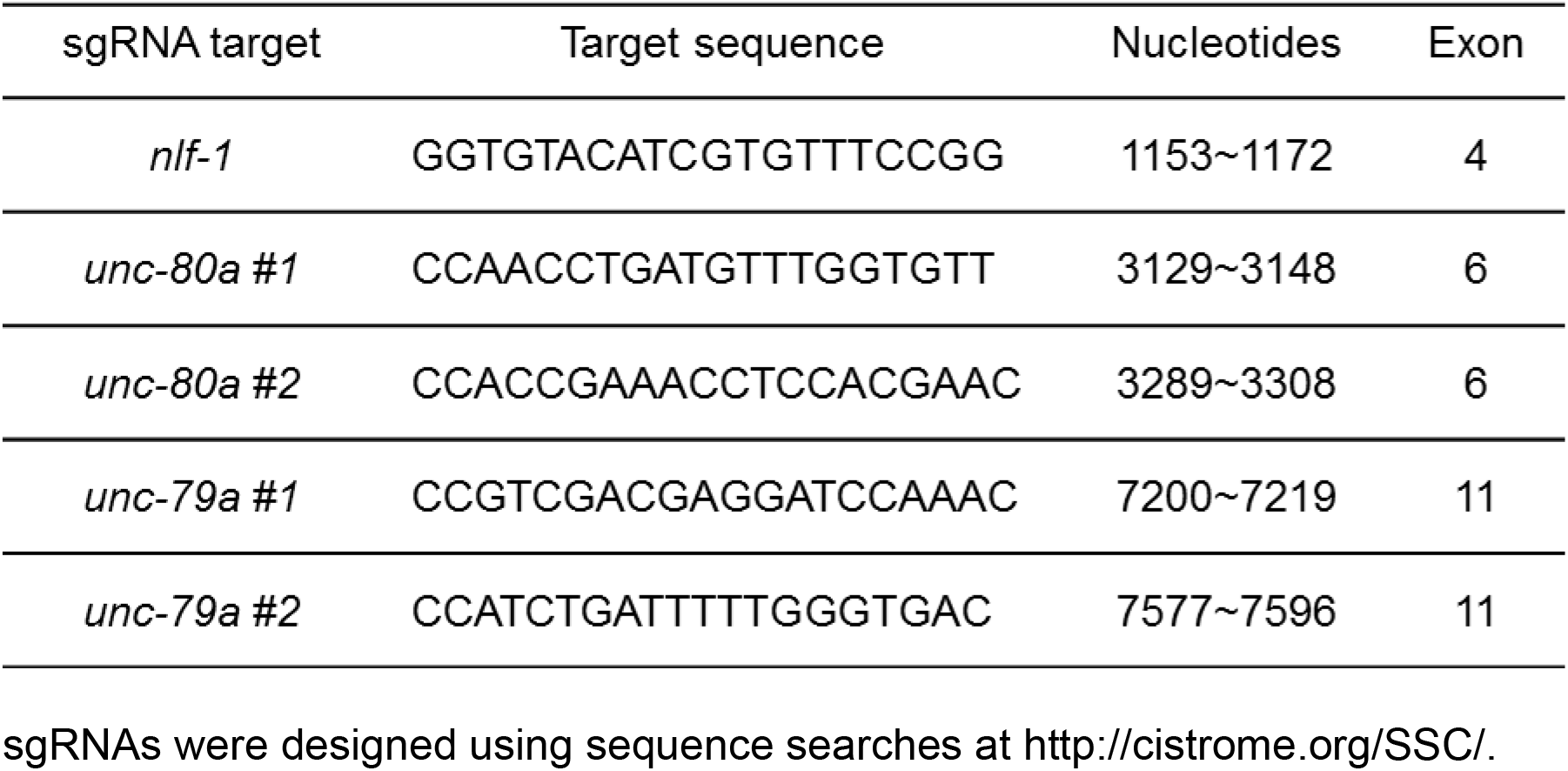
sgRNA target sequences in *nlf-1, unc-80* and *unc-79* for generating genomic lesions using the CRISPR/Cas9 method.

**Table S5.**
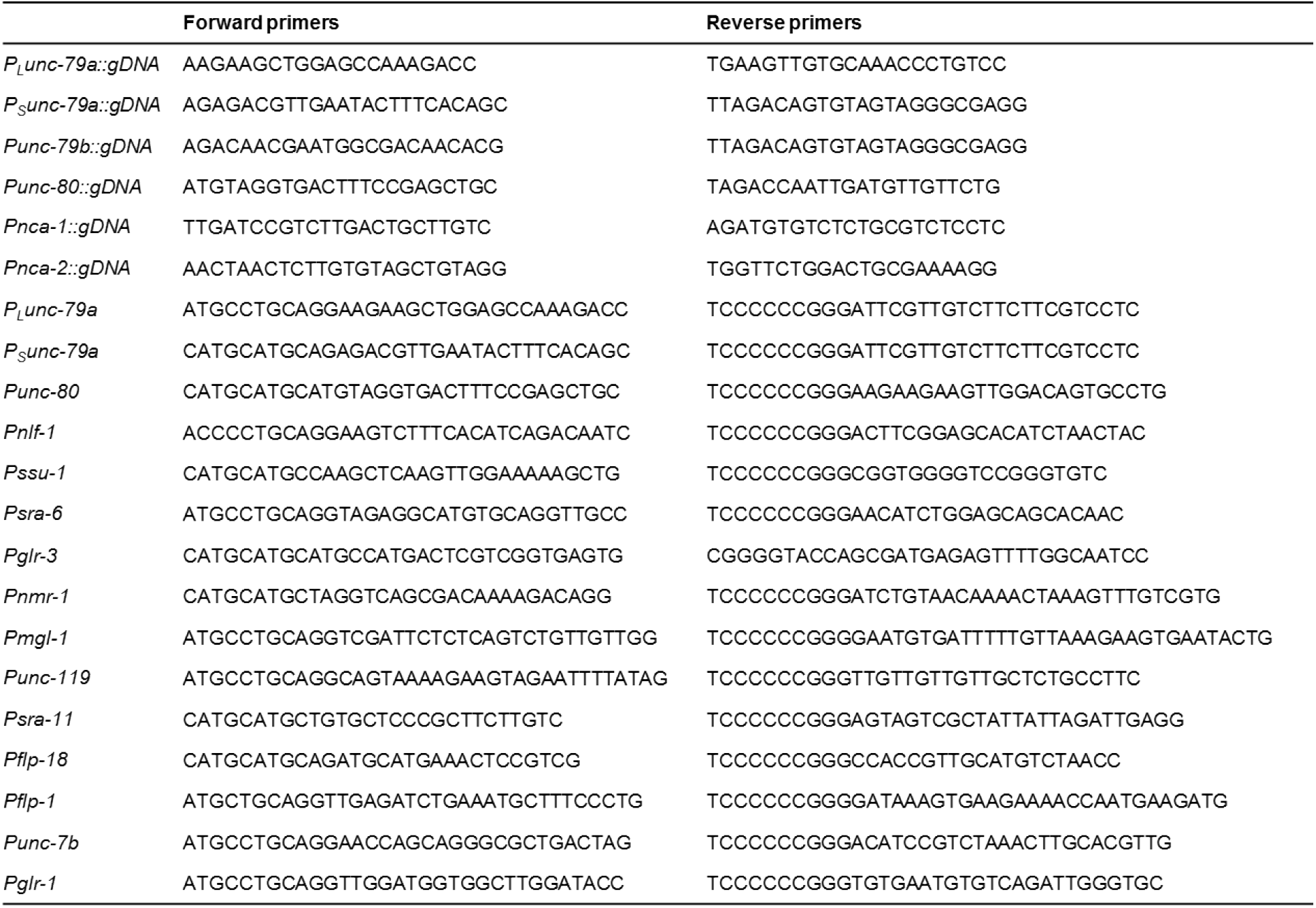
List of PCR primers for amplifying the indicated fragments.

